# Auditory associative word learning in adults: the effects of musical experience and stimulus ordering

**DOI:** 10.1101/2023.11.29.569217

**Authors:** Samuel H. Cosper, Claudia Männel, Jutta L. Mueller

## Abstract

Evidence for sequential associative word learning in the auditory domain has been identified in infants, but adults have proven to have difficulties in this type of word learning. To investigate the role of auditory expertise and auditory stimuli salience in the association of auditory objects and their labels, we tested in the first experiment auditorily-trained musicians versus athletes as a high-level control group and in the second experiment stimulus ordering, contrasting object-label versus label-object presentation. Learning was evaluated from electrophysiological (EEG) recordings during training and subsequent testing phases, as well as accuracy-judgement responses during test. Results revealed for musicians a late positive component in the EEG during testing, neither N400 nor behavioral effects were found at test, while athletes did not show any effect of learning. Moreover, the object-label-ordering group only exhibited emerging association effects during training, while the label-object-ordering group showed an EEG effect during test as well as above chance accuracy-judgement scores. Thus, our results suggest an advantage of musical expertise and more salient label-object-ordering in auditory associative word learning in adults. Implications with respect to developmental differences in auditory word learning are discussed.

## 1 Introduction

To what extent is word learning influenced by perceptual modality? Traditionally, word learning experiments have focused on labeling visual objects (e.g., Breitenstein et al., 2005; Friedrich & Friederici, 2008, 2011; Horst & Samuelson, 2008; McMurray et al., 2012; Sloutsky et al., 2017; Taxitari et al., 2019; Werker et al., 1998). However, recent research has expanded into word learning for entities from other modalities such as tactile (e.g., Miller et al., 2018; Schmidt et al., 2019), olfactory (e.g., Vanek et al., 2021), gustatory (e.g., Yan et al., 2021), and auditory (Cosper et al., 2020, 2022). This distribution of word-learning research in modality does reflect the distribution of sensory modality attributes in the vocabularies of many languages, where the largest attribution of words is to the visual modality (e.g., Chedid et al., 2019; Chen et al., 2019; Lynott et al., 2020; Miklashevsky, 2018; Speed & Brybaert, 2022; Vergallito et al., 2020). In contrast, the auditory modality occupies a middle position in this distribution in most languages, although there is no universal hierarchy of perceptual modality across different vocabularies (Majid et al., 2018). Regarding auditory word learning, it has been shown in early development that words for environmental sounds can be learned associatively in a similar manner compared to words for visual objects (Cosper et al., 2020). Thus, infants are able build associative links between a word (or pseudoword) and an auditory object (e.g., an environmental sound) over time (e.g., McMurray et al., 2012; Regier, 2005; Sloutsky et al., 2017). While adults can also utilize other mechanisms to achieve word learning, such as inferential learning by means of hypothesis testing (cf. Sloutsky et al., 2017; Xu & Tenenbaum, 2007), associative learning remains vital to word learning across the lifespan (Dittinger et al., 2016, 2019; Regier, 2005; Sloutsky et al., 2017). First evidence of associative auditory word learning for adults, however, suggested that their learning of words for sounds is more limited than in children (Cosper et al., 2020, 2022). How strong this limitation is and by what learning factors it might be overcome is yet to be investigated.

The present word-learning study in adults asked whether the above described modality-related constraint on learning can be overcome by external and internal learning factors. Specifically, we aimed to investigate whether behavioral and electrophysiological (EEG) correlates of auditory object–label association can be modulated by i) expertise in the auditory domain and by ii) enhancing the saliency of associated stimuli through the order of presentation. Expertise in the auditory domain will be operationalized by musicianship and contrasted against a high-level control group of athletes. Saliency of the associated stimuli will be operationalized by reversing the order of stimulus presentation compared to our previous word-learning studies with object-label ordering in infants and adults (Cosper et al., 2020, 2022), such that the more salient auditory labels will be presented first across learning trials.

In terms of the cognitive benefits of musical training, musicians have been generally shown to outperform non-musicians in visual, phonological, and executive memory tasks as well as a faster updating of working memory (George & Coch, 2011). Regarding expertise in the auditory modality, musicians have been shown to possess a superior auditory short-term memory as compared to non-musicians (Chan et al., 1998; Cohen et al., 2011; Ho et al., 2003). In an analysis of auditory statistical learning, musicians also showed superior abilities (Pesnot Lerousseau & Schön, 2021) and exhibited enhanced global efficiency and density of the cortical network involved in the learning of tonal sequences as compared to non-musicians (Paraskevopoulos et al., 2017). Moreover, there is evidence that musical expertise and its enhanced cognitive attributes may even transfer to language learning (Schön & Francois, 2011). Both adult musicians and musically-trained children showed improved word learning (visual objects labeled with spoken pseudowords) as compared to non-musicians (Dittinger et al., 2016, 2017).

The decision of including athletes as a control group in the current study is based on the fact that they are also a high-expertise population, yet unrelated to auditory skills (for a similar procedure, see Dittinger et al., 2016). Interestingly, athletes demonstrate positive influences of embodied sport practices in a range of various cognitive tasks, such as visuo-spatial attention processing, and higher level cognitive tasks, such as executive control (Alves et al., 2013). Additionally, athletes exhibit advantages in attentional cuing, processing speed, and varied attention paradigms as compared to non-athletes or non-professional athletes (Voss et al., 2010). Implicit memory advantages have also been found for experienced athletes, although these results are limited to the context of the sport (e.g., soccer players in soccer-specific situations; Zoudji & Thon, 2003). Furthermore, elite-athletes exhibited an advantage over recreational-athletes in sustained visual attention, but again these advantages are not extended to non-sports-specific cognitive measures, such as verbal memory span (Heppe et al., 2016).

In addition to auditory expertise, we also investigated the effect of stimulus saliency by manipulating the order in which stimuli are presented. This is particularly relevant for auditory associative word learning, as auditory information is by nature fleeting and does not persist over time. While environmental sounds and speech are processed in similar ways in principle, for example, at the level of spectrotemporal analysis (Klein et al., 2003), but also with respect to processing of context and meaning (Ballas & Howard, 1987; van Petten & Rheinfelder, 1995), speech is more salient. As indicated by activation patterns in the auditory cortex and in behavioral analyses, discrimination of speech is better than discrimination of environmental sounds in natural auditory scenes containing both stimuli (Renvall et al., 2021). However, neural effects of conceptual priming (N400) have indicated that familiar environmental sounds may elicit an earlier onset of the N400 than those elicited by words, although conceptual processing of verbal and non-verbal stimuli is considered to be similar in nature (Orgs et al., 2006). Behavioral measures have also shown increased accuracy rates in the processing of auditory-verbal information in relation to auditory-nonverbal information both in the processing of visual objects pairings with either linguistic or non-linguistic stimuli (Calignano et al., 2021) and in statistical learning (Lukics & Lukács, 2022; Thiessen, 2010). Short-term memory accuracy is also higher in tasks with verbal stimuli than with nonverbal stimuli in both the auditory and the visual modalities (Talamini et al., 2022). Taken together, these studies suggest a processing and memory advantage for speech compared to non-speech stimuli. Yet, when sequences of speech and non-speech stimuli are presented together, not only does the salience of each single stimulus play a role, but also how the two interact with each other. We know that both words and environmental sounds can be used as cues to meaning in principle (Orgs et al., 2006; van Petten & Rheinfelder, 1995), but we do not know how the factors detailed above, saliency and memorizability, may impact on learning if auditory objects and spoken labels are presented in sequence.

The current study uses an adaptation of the word learning paradigm of Friedrich and Friederici (2008, 2011), with a combination of behavioral and electrophysiological measurements to allow for an in depth analysis of associative word learning in adults. In two experiments, auditory objects (environmental sounds) and pseudowords (labels) are presented sequentially in pairs across learning and testing phases, while an EEG is recorded and accuracy ratings are obtained. First, two expert populations are tested, musicians and athletes, and second, a general-population group is tested in a label-object-ordering experiment and compared to a previously reported study using object-label ordering (Cosper et al. 2022). For the musicians, we predicted the group would show behavioral (i.e., above-chance accuracy on a judgement task) and electrophysiological evidence of word learning in the testing phase (i.e., N400 effect). For the athletes, we predicted this group would be similar to the auditory-sequential condition of Cosper and colleagues (2022) and not show learning effects. For the second experiment, with reversed label-object ordering, we expected learning effects due to an attentional and memory advantage of labels that should facilitate linking related stimuli in the learning task, as evidenced in above-chance accuracy in the judgment task as well as an N400 effect in the EEG.

## 2 Experiment 1: Musicians and Athletes

### 2.1 Materials and methods

#### 2.1.1 Participants

A total of 48 participants (24 females and 24 males) with ages ranging from 19 to 29 years of age (*M* = 23.32 years; *SD* = 2.84 years) in two different groups participated in the study: musicians and athletes. All participants were found through the University of Osnabrück, native German speakers, right-handed, reported normal hearing and normal/corrected eye-sight, and did not suffer from any neurological conditions (including learning impairments, ADHD, or dyslexia). In order to be included in the final dataset, each participant must have retained at least 75% of each trial type after artifact rejection and ICA correction, which is described in more detail below.

For the musicians, a group of 24 students participated, one participant was not a student but had 10 years of experience playing in a professional symphony. One participant (*n*=1) was excluded, as inclusion criteria after pre-processing was not met. The final dataset for musicians included 23 individuals (11 female), aged 19 to 29 (*M* = 24.02 years; *SD* = 5.57 years). In order to be included in the study, musicians had to have been playing an instrument for at least 11 years (for similar requirements, see Dittinger et al., 2016) as well as being enrolled as a music student in a Bachelor or Master program at the University of Osnabrück (see above for the one exception). The musicians reported having played an instrument for an average of 15.04 years (*SD* = 2.59 years). Instruments reported included 10 pianos, seven violins, two trumpets, one drums, one horn, one cello, and one trombone.

For the athletes, 24 students (12 female) participated with ages ranging from 19 to 29 years of age (*M* = 22.74 years; *SD* = 2.68 years). In order to be included in the study, athletes needed to be active in sports with regular training sessions (at least 3.5 hours a week) for at least the past five years. Athletes reported participating in competition sports ranging between 5 to 19 years (*M* = 10.33 years; *SD* = 3.59 years), and training from 3.5 to 7 hours a week (*M* = 4.84 hours per week; *SD* = 1.01 hours). Sport disciplines were reported in groups: team sports (soccer/football, handball, volleyball, water polo, hockey, and table tennis; *n* = 14), individual sports (swimming, track and field, triathlon, bouldering; *n* = 7), and martial arts (judo, kickboxing, and karate; *n* = 3). For an overview of all participants, see Table 1.

**Table 1:** Overview of participants by sample size and age per group along with average time played (instrument for musicians, competition sports for athletes, and non-applicable for the object-label-ordering group and the label-object-ordering group) and average number of trials per condition included in the final dataset per participant group. Standard deviation was also indicated for all averages. The participant information from the object-label-ordering group (marked *) was from a previous publication (auditory sequential; Cosper et al., 2022).

| Participant Group | Sample Size | Average Age (SD) | Average years trained (SD) | 1<sup>st</sup> Half Training – Consistent (SD) | 1<sup>st</sup> Half Training – Inconsistent (SD) | 2<sup>nd</sup> Half Training – Consistent (SD) | 2<sup>nd</sup> Half Training – Inconsistent (SD) | Testing – Matching (SD) | Testing – Violated (SD) |
| --- | --- | --- | --- | --- | --- | --- | --- | --- | --- |
| Musicians | 23 | 24.02 years<br>(5.57 years) | 15.04 years<br>(2.59 years) | 30.8 trials<br>(1.5 trials) | 30.7 trials<br>(1.9 trials) | 30.9 trials<br>(1.3 trials) | 30.6 trials<br>(1.5 trials) | 14.9 trials<br>(1.1 trials) | 14.8 trials<br>(1 trial) |
| Athletes | 24 | 22.74 years<br>(2.68 years) | 10.33 years<br>(3.59 years) | 30.3 trials<br>(1.9 trials) | 30.2 trials<br>(1.9 trials) | 29 trials<br>(3.1 trials) | 28.5 trials<br>(4.6 trials) | 13.8 trials<br>(2.5 trials) | 14 trials<br>(2.2 trials) |
| Object-Label-Ordering* | 22 | 23.86 years<br>(3.4 years) | - | 31.2 trials<br>(1.2 trials) | 31.2 trials<br>(1.2 trials) | 30.4 trials<br>(1.6 trials) | 30.6 trials<br>(0.7 trials) | 15.5 trials<br>(0.7 trials) | 15.1 trials<br>(1 trial) |
| Label-Object-Ordering | 23 | 23.36 years<br>(5.23 years) | - | 29.9 trials<br>(3.1 trials) | 30.1 trials<br>(2.2 trials) | 29.2 trials<br>(3.1 trials) | 29.6 trials<br>(2.3 trials) | 13.7 trials<br>(2.2 trials) | 14.1 trials<br>(1.9 trials) |

None of the participants of this experiment had previously participated in other auditory associative word learning experiments. Participants of all groups gave written informed consent before participation. All participants were compensated for their time with either credit towards their degree program (mandatory participation in experiments for Psychology and Cognitive Science Bachelor programs) or 10 Euro. The experiment was approved by the ethics committee of the University of Osnabrück and conformed to all aspects of the Declaration of Helsinki (World Medical Association, 2013).

#### 2.1.2 Stimuli

The stimuli utilized in the current experiment contained 16 environmental sounds from the NESSTI database for environmental sounds (Hocking et al., 2013), cut to 950 ms in length, and 16 pseudowords derived from existing German nouns with sound files 750 ms in length. All stimuli were previously used in auditory associative word learning experiments in both infants and adults (Cosper et al., 2020, 2022). All files were saved as monaural .WAV files, digitized at 44100 Hz, and presented over two speaker channels at a comfortable volume. The experiment was presented using the Presentation software (Neurobehavioral Systems, Inc., Berkeley, CA, USA, version 20.0).

#### 2.1.3 Procedure

For the experiment, participants were seated in front of a computer monitor, outfitted with speakers on either side. Buttons for the behavioral analysis were on the table within reach. Participants were instructed to pay attention to the sounds and words presented, as they would be asked questions about them at a later point, while looking at the screen. Ultimately, the participants were ignorant to the goal of the study and were not explicitly instructed to attend the sound-pseudoword (or pseudoword-sound) pairings and their consistency. The experiment was split into two sections: a training phase and a testing phase. The training phase consisted of 128 trials in which environmental sounds and pseudoword combinations were presented. Of the 16 environmental sounds, eight were consistently presented with a single pseudoword eight times each (for a total of 64 trials). The remaining eight environmental sounds and eight pseudowords were paired with one another only once, creating 64 unique combinations in as many trials (for similar training phases in associative word learning, see Cosper et al., 2020, 2022; Friedrich & Friederici, 2008, 2011). Inconsistent pairings were included to test online association during the learning phase. The testing phase utilized only the environmental sounds and pseudowords of the consistent pairings in the training phase, presented either matching (identical to the consistent pairing in training) or violated (new combinations of sounds and pseudowords, violating built pairings in training). Each pairing was presented twice for a total of 16 pairings in each condition and 32 trails overall. Figure 1A provides an example of the sound-pseudoword combinations in the training phase and Figure 1B gives an example of the testing phase. Eight randomized lists were created to counterbalance stimuli across training conditions, redistribute pairings, and to invert presentation lists, in order to test overall associative word learning abilities and not individual stimuli pairing preferences. A fixation cross was always presented on the screen.

**Figure 1:**
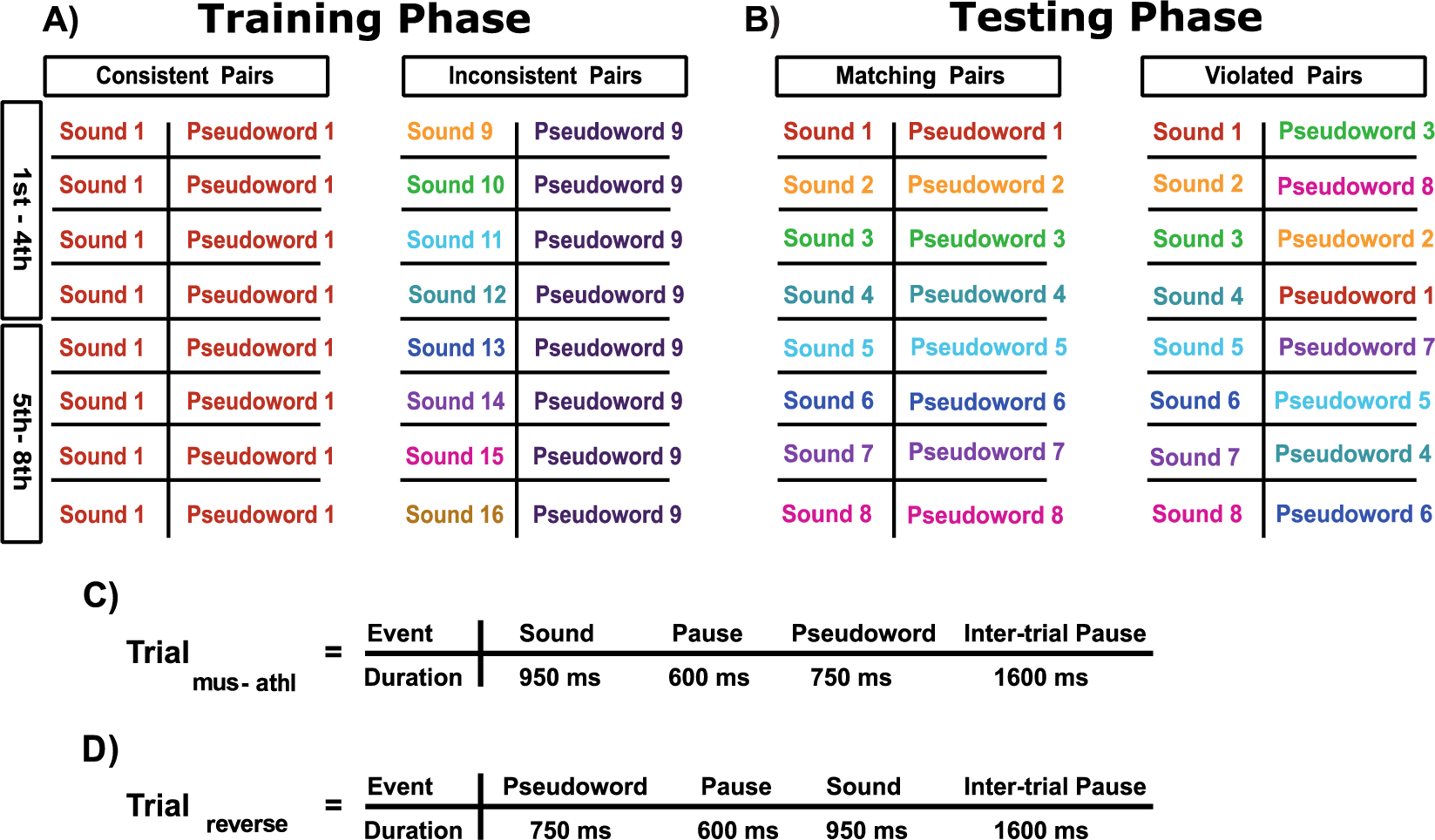
Experimental setup. (A) Example of pairings of sound-pseudoword combinations for the training phase as presented to the musician and athlete groups. (B) Example of pairings of sound-pseudoword combinations for the testing phase as presented to the musician and athlete groups. (C) Trial structure for the musician and athlete groups. (D) Trial structure for the label-object-ordering group. Colors used for visual emphasis. Figure adapted from Cosper and colleagues (2022).

Each trial was structured identically across the experiment. For experiment 1, sounds were presented first, then a 600 ms inter-stimulus pause, followed by the pseudoword (see Figure 1C). After the final stimulus in the pairing was presented, a 1600 ms inter-trial pause was presented. Between the training and testing phase, participants has a 1-2-minute break, in which the instructions for the testing phase were also given. In the testing phase, participants were additionally asked to provide behavioral input after the presentation of each pairing, indicating if the pairing was correct by means of button pressing. The assignment of left and right buttons (for yes and no answers) was counterbalanced across the randomization lists. The experiment lasted 14 minutes in total.

#### 2.1.4 EEG Processing

All data were collected at the Kindersprachlabor of the University of Osnabrück. EEG signals were measured with a REFA amplifier (Twente Medical Systems International, Oldenzaal, The Netherlands) using Ag/AgCl electrodes at extended standard 10-20 electrode positions in 64-channel TMSi caps (Twente Medical Systems International, Oldenzaal, The Netherlands). Eye movement-related activities were measured by monopolar electrodes placed above and below the left eye and on either temple. Impedances were kept below 5 kΩ. The EEG data were recorded continuously using the TMSi Polybench software (Twente Medical Systems International, Oldenzaal, The Netherlands) at a sampling rate of 512 Hz, average reference, and a ground electrode located on the left collarbone.

Pre-processing of EEG data was done in MATLAB (The Mathworks Inc., Natick, MA, USA, version 2022b) using EEGLAB (Delorme & Makeig, 2004, version 14_1_1b) following the same protocol as Cosper and colleagues (2022). First, a high-pass filter of 1 Hz (-3dB, cutoff frequency of 1.38 Hz) and a low-pass filter of 30 Hz (-3dB, cutoff frequency of 31.16 Hz) were applied to the continuous EEG data. The data was then epochized, with epochs time-locked to the onset of the second stimulus in the presented pairing with a length of 1200 ms and a pre-stimulus baseline of 200 ms. For clarity, the analyses for musician, athlete, and object-label- ordering groups were conducted on the pseudoword and the analyses for the label-object-ordering group were conducted on the environmental sound. Subsequently, EEG data were re-referenced to linked mastoids. Artifact rejection was conducted by means of manual inspection. Following, a semi-automatic independent component analysis (ICA) was applied to each dataset in order to correct eye-movement artifacts. Afterwards, the ICA weights of the 1 – 30 Hz filtered dataset were applied to a dataset with a high-pass filter of 0.3 Hz (-3dB, cutoff frequency of 0.36 Hz) and a low-pass filter of 30 Hz (-3dB, cutoff frequency of 31.16 Hz). All statistical analyses were conducted using this 0.3 – 30 Hz dataset. As mentioned above, participants must have met the inclusion criteria to remain in the final dataset. The average number of trials and their standard deviations for each participant group can be seen in Table 1.

#### 2.1.5 Statistical Analysis

The behavioral results are presented as the accuracy of participants’ responses and was assessed in R (R Core Team, 2020) using a one-sample t-test against chance, µ = 0.5.

EEG data were statistically analyzed for each participant group in MATLAB (The Mathworks Inc., Natick, MA, USA) using the FieldTrip toolbox (Oostenveld et al., 2011, version 20220104). Cluster-based permutation analyses were conducted by means of dependent samples t-tests for each sampling point with a set alpha threshold of 0.05. A minimum number of neighborhood channels was set to 2 for a significant sample to be included in the clustering algorithm. A permutation test, using the Monte Carlo method (Maris & Oostenveld, 2007), was applied with 1000 randomizations and an alpha threshold of 0.05. Reference electrodes and EOG electrodes were excluded from the cluster-based permutation test. Two time windows (TWs) were selected for the analyses. An N400 TW was selected from 400 – 800 ms post-stimulus, pre-selected in order to compare to previous auditory associative word learning experiments (Cosper et al., 2022). Additionally, a later TW was selected to assess possible later effects: 800 – 1200 ms.

The EEG statistical analyses were conducted in three comparisons: first half of the training phase, second half of the training phase, and the testing phase. For the first half of the training phase, cluster-based permutation tests were applied to test differences between consistent pairings versus inconsistent pairings for the first through fourth presentations of each target stimulus (pseudoword or environmental sound). For the second half of the training phase, cluster-based permutation tests were applied to the fifth through eighth presentations of each target stimulus to assess differences between consistent and inconsistent pairings. Violated and matching pairings were assessed in the testing phase for each group. It is important to note that as cluster-based permutation tests are applied here, TWs reported in the results section are not meant to be determinations of absolute TWs of effects nor are cluster-based permutation tests suitable for the determination of effect latency (cf. Sassenhagen & Draschkow, 2019). Thus, results presented below are approximate effect TWs and only describe clusters of electrodes in which significant differences are found.

### 2.2 Results

#### 2.2.1 Behavioral Results

The behavioral analysis of the musician group showed that participants correctly identified correct and incorrect sound-pseudowords pairings in the testing phase only 52% of the time (*M* = 16.74 correct responses; *SD* = 2.59 responses). A one-sample t-test indicated that the musicians were not above chance identifying matching and violated pairings, *t*(22) = 1.3379, *p* = 0.195 (see Figure 2).

**Figure 2:**
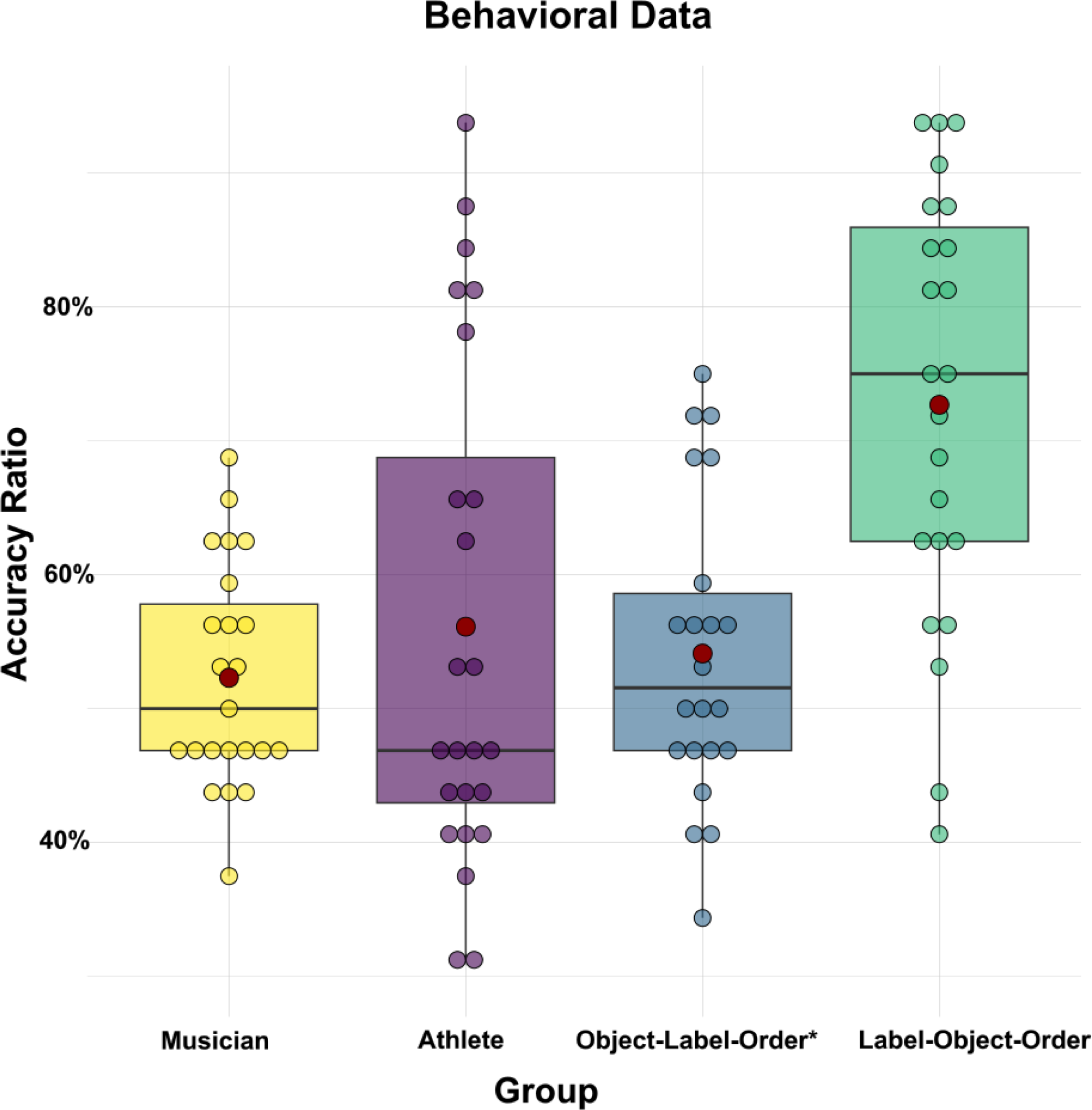
Behavioral results of the testing phase. The distribution of the accuracy ratios for each participant is presented per group. Chance level is at 50%. The boxes represent the upper and lower quartile, while the whiskers represent variability beyond the quartiles. The line within the boxes represents the median accuracy ratio, the red dot represents the mean, and the colored dots represent the individual accuracy ratios of the participants. The data for the Object-Label-Ordering group is from a previous publication (auditory sequential; Cosper et al., 2022).

The behavioral analysis of the athletes group yielded a 56% accuracy ratio (*M* = 17.96 correct responses; *SD* = 5.95 responses). A one-sample t-test indicated that athletes were not above chance in correctly identifying matching and violated sound-pseudoword pairings in the testing phase, *t*(23) = 1.5791, *p* = 0.128 (see Figure 2).

#### 2.2.2 EEG: Training Phase

For the musician group, no clusters with significant processing differences between inconsistent over consistent sound-pseudoword pairings in the first or the second half of training were found in either the 400 – 800 ms or the 800 – 1200 ms TW (see Figure 3A).

**Figure 3:**
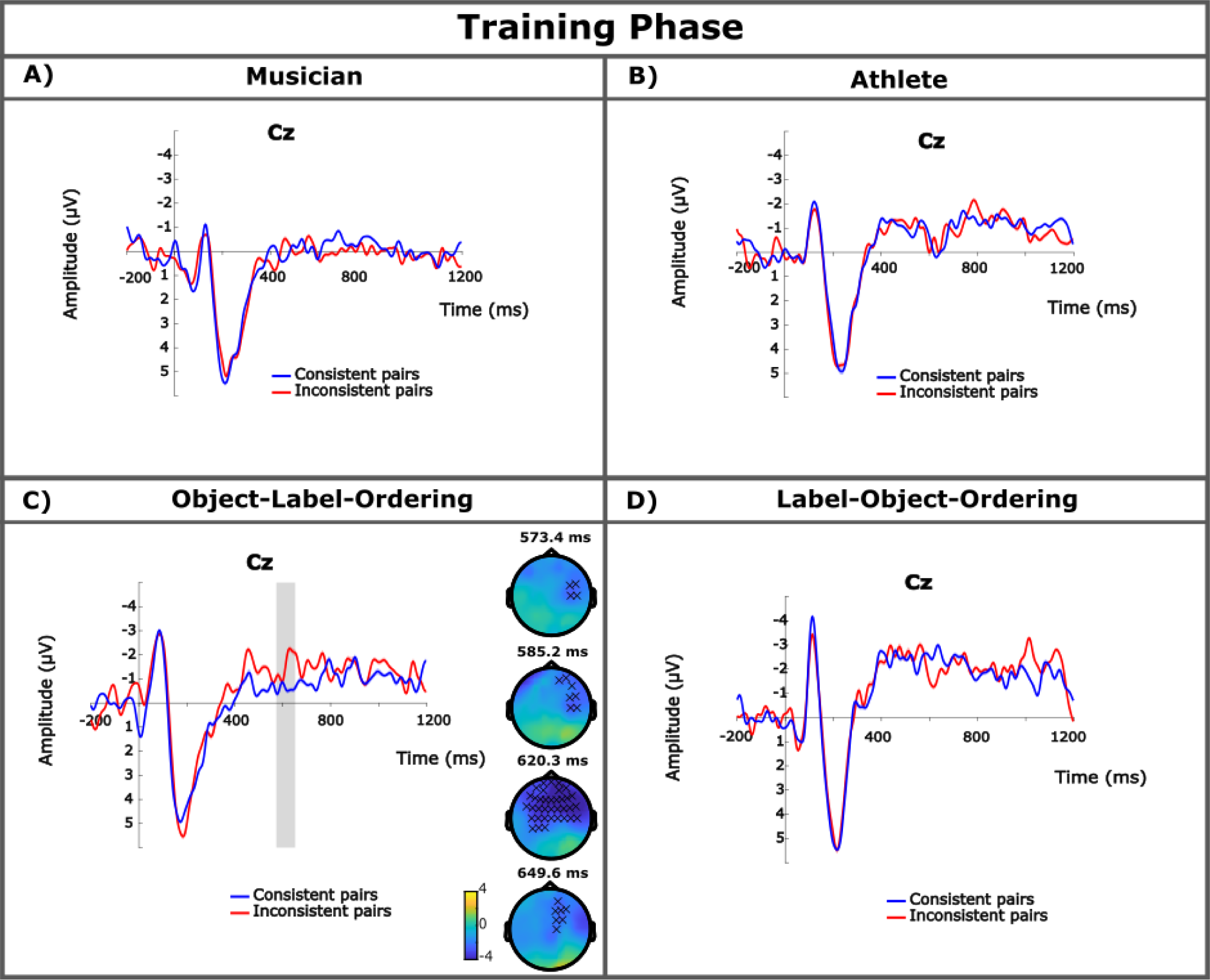
Training phase results. ERPs are presented time-locked to the onset of the second stimuli in the pairing for the second half of the training phase of each experiment (A, B, C represent pseudowords, D represents environmental sounds). The Cz electrode was selected as a representative electrode for all of the experiments. The blue lines indicate the consistent pairs while the red lines indicate the inconsistent pairs. The lighter-shaded lines indicate the standard error of the waveform. Gray bars are used to illustrate the time window where significant differences were found between the conditions. Topological difference maps from the cluster-based permutation test are presented only where significant differences were found and show the electrode distribution in the cluster over time (the X symbol indicates *p* < 0.05). Results of Object-Label-Ordering (C) are reprinted from a previous publication (Cosper et al., 2022).

In the athlete group, no clusters showing significant processing differences between inconsistent and consistent sound-pseudoword pairings were found in the first or the second half of the training phase in either the N400 or late TW (see Figure 3B).

#### 2.2.3 EEG: Testing Phase

The cluster-based permutation in the testing phase of the musician group yielded a single positive cluster with a significant processing difference between violated and matching sound-pseudowords (*p* = 0.014). This difference was found between 406 – 512 ms after the onset of the pseudoword with a broad topological distribution during peak cluster activation. No clusters of significant differences were found for the later TW (see Figure 4A).

**Figure 4:**
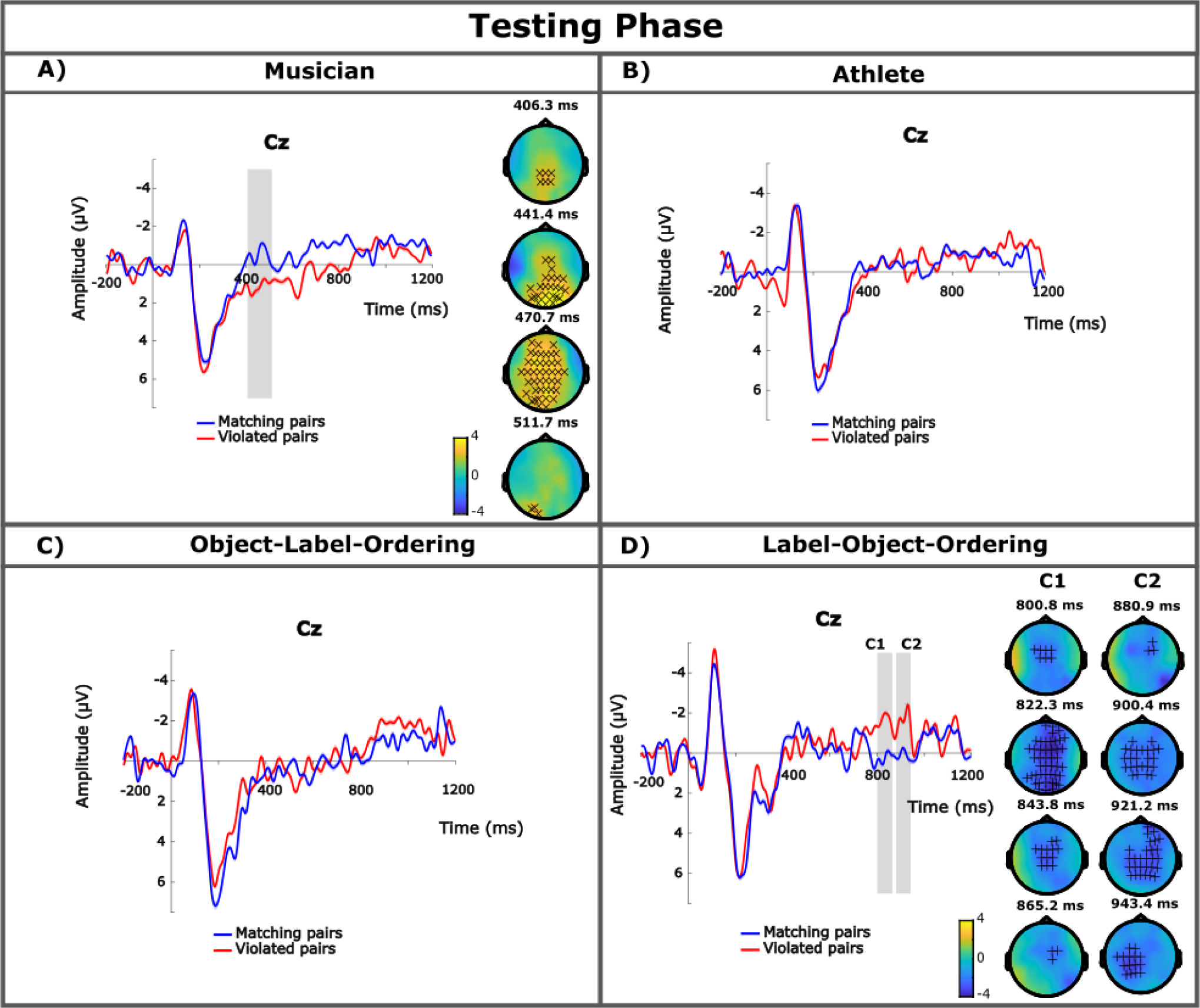
Testing phase results. ERPs are presented time-locked to the onset of the second stimuli in the pairing for the testing phase of each experiment (A, B, C represent pseudowords, D represents environmental sounds). The Cz electrode was selected as a representative electrode for all of the experiments. The blue lines indicate the matching pairs while the red lines indicate the violated pairs. The lighter-shaded lines indicate the standard error of the waveform. Gray bars are used to illustrate the time window where significant differences were found between the conditions. In the case of multiple clusters, the clusters are labeled C1 and C2. Topological difference maps from the cluster-based permutation test are presented only where significant differences (or trends) were found and show the electrode distribution in the cluster over time (the + symbol indicates 0.1 > p > 0.5 and the X symbol indicates *p* < 0.05). Results of Object-Label-Ordering (C) are reprinted from a previous publication (Cosper et al., 2022).

For the athlete group, no clusters of significant processing differences were found between the violated and matching sound-pseudoword pairs in the testing phase for either the N400 or the later TW (see Figure 4B).

## 3 Experiment 2: object-label-ordering and label-object-ordering

### 3.1 Materials and methods

#### 3.1.1 Participants

Participants in the object-label-ordering group were 22 students from a previous experiment (auditory sequential; Cosper et al., 2022). All participants (16 female) were included in the final dataset, aged 19 to 32 years of age (*M* = 23.86 years; *SD* = 3.4 years). Table 1 provides an overview of all participants.

For the label-object-ordering group, 24 students were recruited to participate in the study. One participant (*n* = 1) was removed for not meeting inclusion criteria, leaving 23 students (11 female) in the final dataset, aged 18 to 29 years of age (*M* = 23.36 years; *SD* = 5.23 years). For an overview of all participants, see Table 1. None of the participants of this experiment had participated in any other auditory associative word learning experiment. The recruitment of participants as well as all inclusion and exclusion criteria were identical to experiment 1.

#### 3.1.2 Stimuli and Procedure

The procedure of experiment 2 followed the procedure of experiment 1. Note, the example stimuli combinations in Figure 1A and 1B are provided for experiment 1 as well as for the object-label-ordering group of this experiment. The parings for the label-object-ordering group were presented in the opposite order. For the object-label-ordering group, environmental sounds were first presented, followed by an inter-stimulus pause of 600 ms, then a pseudoword was presented (identical to experiment one; see Figure 1C). For the label-object-ordering group, pseudowords were presented first, then a 600 ms inter-stimulus pause, followed by the environmental sound (see Figure 1D). The data for the object-label-ordering group is from a previous publication (auditory sequential; Cosper et al., 2022) reporting the same findings for means of comparison; however, the data have been newly analyzed for the later time window (800 – 1200 ms) in both the training and testing phases to be comparable to the other groups in experiment 1 and 2.

### 3.2 Results

#### 3.2.1 Behavioral Results

The object-label-ordering group of the previous study did not indicate any behavioral results above chance (auditory sequential; Cosper et al., 2022). For the label-object-ordering group, a one-sample t-test indicated that the participants were significantly above chance in identifying matching and violated pseudoword-sound pairings in the testing phase at 73% (*M* = 23.36 correct responses; *SD* = 5.03 responses), *t*(22) = 6.7677, *p* < 0.001 (see Figure 2).

#### 3.2.2 EEG: Training Phase

The previous study indicated a single cluster of significant differences in the second half of the training phase for the object-label-ordering group (auditory sequential; Cosper et al., 2022). No clusters of significant processing differences were found in the later TW in either the first or second half of training (see Figure 3C).

The analysis of both the first and second half of training for the label-object-ordering group yielded no clusters in which significant differences between inconsistent and consistent pseudoword-sound pairings were found in either TW (see Figure 3D).

#### 3.2.3 EEG: Testing Phase

As previously reported, the object-label-ordering group, no clusters in which significant processing differences between matching and violated sound-pseudoword pairs were found in either the N400 TW (auditory sequential; Cosper et al., 2022). Further analysis similarly found no clusters of significant processing differences in the later TW (see Figure 4C).

The analysis of the label-object-ordering group did not yield any clusters in which significant differences were found between violated and matching pseudoword-sound pairs in either the 400 – 800 ms or the 800 – 1200 ms TW. However, two trends were identified in the later TW. Cluster 1, from 801 – 865 ms after the onset of the sound, narrowly escaped significance differences between processing of violated and matching pairings (*p* = 0.068). Cluster 2, from 881 – 943 ms, also yielded a trend in the differences between the processing of violated over matching pairs (*p* = 0.074). These clusters in which a trend has been identified between the conditions in the testing phase can be seen in Figure 4D.

## 4 Discussion

To improve our understanding of external and internal modulators of associative auditory word learning in adults we conducted two behavioral and ERP experiments in which novel word forms were sequentially linked to novel environmental sounds. First, two expert groups were tested on object-label ordered learning sequences. Second, a non-expert group was tested on label-object ordered sequences and compared to a non-expert group previously tested on object-label ordered sequences (Cosper et al., 2022). For musicians, electrophysiological analysis of the training phase did not reveal an N400 effect, and at test, behavioral response accuracy was not above chance. However, we observed a late EEG effect in the testing phase with more positive-going ERP responses for violated than matching stimulus pairs. In the athlete group, neither behavioral nor ERP analyses of the training and testing phases indicated successful auditory associative word learning. For the two non-expert groups, label-object-ordering revealed an above-chance response accuracy in the testing phase as well as a marginally significant late negativity for violated than matching pairs in the testing phase. In contrast object-label-ordering did not indicate above-chance response accuracy or yield any ERP effects during the testing phase (cf. Cosper et al., 2022). These results thus suggest that both experimentally manipulated learning factors, auditory expertise and auditory saliency from stimulus ordering, impacted on the learnability of stimulus pairs. We will discuss the potential impact of these factors on the behavioral and electrophysiological outcome of auditory associative learning in more detail below.

For the testing phase, musicians showed a positive cluster of electrodes for the processing of violated over matching sound-pseudoword pairs. Although the latency of this effect generally aligns with the expectation of an N400 (Bechtold et al., 2023; Cosper et al., 2022; Kutas & Federmeier, 2000, 2011), given absolute latency and time window limitations of cluster-based permutation analyses (Sassenhagen & Draschkow, 2019), the polarity is reversed and does not mirror the polarity for the elicited component to be an N400. We suggest that the elicited effect is a reflection of the late positive component (LPC), which is typically found in musicians as a response to incongruous elements in musical melody and rhythm, increasing with familiarity and musical expertise (cf. Besson & Faïta, 1995; Calma-Roddin & Drury, 2020; Featherstone et al., 2013; Miranda & Ullman, 2007). In their analysis, Featherstone and colleagues (2013) reported an LPC (also related to the P600 component for syntactic processing) in musicians between 500 – 700 ms after the onset of the target stimulus. The authors, moreover, reported that non-musicians exhibited an N500 response (related to the N400 for semantic processing) for the same stimuli presentations. Similarly, Besson and Faïta (1995) reported an LPC (400 – 800 ms) for congruity with larger amplitude for musicians than for non-musicians. The interpretation of these findings is that individuals with musical expertise (i.e., musicians) process incongruent semantic relations in musical stimuli differently than non-musicians and that the neural underpinnings of these processes may change with expertise (Besson, 1998; Besson et al., 1994; Featherstone et al., 2013; Miranda & Ullman, 2007; Steinbeis et al., 2006). Thus, in our study, it may not only be the case that musicians implicitly learned a link between auditory objects and labels, but musicians may also have processed these associations differently than non-musicians. As no behavioral learning effects were found, we interpret the LPC effect as an indication of implicit learning that does not be necessarily need to be lexical-semantic in nature. As N400 effects have been found with musicians in associative word learning with visual objects (e.g., Dittinger et al., 2016, 2017, 2019), it is unlikely that the LPC simply reflects word learning in musicians. Instead, the LPC may simply reflect a process akin to auditory statistical learning. This suggestion is supported by experiments in which two auditory elements were sequentially linked, and in which similar positivities were found (e.g., Friederici et al., 2011; Mandikal Vasuki et al., 2017). Thus, the present results are consistent with previous findings in musicians evidencing enhanced auditory processing and memory (Chan et al., 1998; Cohen et al., 2011; George & Coch, 2011; Ho et al., 2003; Paraskevopoulos et al., 2017), and the transfer of musical expertise to language processing (Dittinger et al., 2016, 2017, 2019; Pesnot Lerousseau & Schön, 2021; Schön & Francois, 2011).

In contrast to the musicians, no significant effects were found in the athlete group in either behavioral or ERP analyses. Athletes were not above chance levels at identifying correct and incorrect sound-pseudoword pairings. Furthermore, electrophysiological brain signals did not indicate any significant processing differences between matching and violated sound-pseudoword pairings during the testing phase. On the one hand, these findings corroborate and replicate the findings of Cosper and colleagues (2022) showing that adults (without auditory expertise) are not able to map spoken labels onto auditory environmental sounds in sequential object-label ordered presentation. On the other hand, the absence of findings for the athlete group also support the findings of Heppe and colleagues (2016) as well as Zoudji and Thon (2003) in that the attentional and cognitive advantages of athletes do not transfer beyond sport-related topics.

An important outcome of the present study was the non-replication of the N400 effect in the training phase that had been reported in an adult study of word learning by Cosper and colleagues (2022). In this previous study, the N400 effect in the training phase was interpreted as being related to the sequentiality of the stimulus presentation, whereas sequential presentation yielded an N400 response in training and simultaneous presentation did not yield an N400 response. The present results from the three ERP experiments including the object-label-order condition (auditory sequential; Cosper et al., 2022) suggest that our previous interpretation might have been inadequate. The non-replication of the N400 effect could be either the result of insufficient power (yet participant numbers were comparable) or hinting to the possibility that there are other factors influencing the presence of the N400 effect during associative word learning which is expected to occur only in the second half of the training phase. Such factors could be a high sensitivity to the dynamics of learning (for example, if sounds and words are linked only in the very last repetitions, the effect would be not observable) or subtle differences in motivation and attention on the side of the participants. In order to dissect the influences behind the N400 during learning, further experiments are needed and should be the focus of future research.

Regarding stimulus order, the behavioral analysis of the testing phase yielded above-chance accuracies only for the label-object ordered but not for the reverse condition. This suggests that adults are better learners in in auditory-sequential associative word learning, when the more salient auditory label appears first (cf. Cosper et al., 2022). Despite above-chance behavioral responses, the ERP analysis of the testing phase for the label-object-ordering group did not meet the threshold of significance for processing differences between matching over violated pseudoword-sound pairings. Two clusters were identified with nearing statistically significant processing differences as identified by an N400-like component. Although these trends appear much later than the target TW of the N400, later N400-like responses have been seen in both children (for a review, see Junge et al., 2021) and in adults (auditory-simultaneous associative word learning; Cosper et al., 2022). Alternatively, the identification of the sounds could have taken longer given that the sounds were not as initially unique in their identification as the pseudowords. Even if sound identification did not always take longer, but might have been more variable, this could have led to the increased latency and non-significance of the effect which may have captured just the final processing stage in which all the unique sounds were identified. Note that it is unlikely that the environmental sounds per se yielded a later effect. Orgs and colleagues (2006) provided evidence that contextual priming between written words and environmental sounds elicited an N400 effect with an even earlier onset than a picture and a written word. However, the environmental sound stimuli used by Orgs and colleagues (2006) were much shorter than the environmental sound stimuli used in the current set of experiments (300 ms versus 950 ms, respectively). It is therefore possible that the associative memory processes, be it from the encoding process of association between the two stimuli presented or from the maintenance of the environmental sound, may have contributed to the delay of the N400 in the object-label-ordering condition. Thus, the late negative component in the label-object-ordering group might similarly reflect similar a violation of lexico-semantic expectation, as suggested in Cosper and colleagues (2022), but this proposal would need to be confirmed in an independent study.

In sum, given the advantages in behavioral response accuracy and the observed trends in the ERP analysis of the testing phase, we suggest that auditory-sequential associative word learning does benefit from label-object-ordering in comparison to object-label-ordering in a general, non-musician population. This may be due to the increased saliency and memorizability of words compared to sounds which may result in a better matching of words and sounds when the sounds are presented in the second position in the sequence (cf. Renvall et al., 2021). In contrast, when sounds are presented first, it may be harder to form a memory representation and thus, when the word appears in the second position, matching may be difficult because the sound has already faded.

We caution to the overgeneralization of our results, as we are limited in our scope of interpretation. Although musicians and athletes were tested as expertise groups, groups assignment was only substantiated by the self-reports of the individual participants and no measures were conducted to identify the expertise of either group. In future studies, this should be addressed, for example, by investigating individual indexes of modality-specific (i.e., visual and auditory) short-term memory recognition accuracies with both verbal and nonverbal stimuli. Additionally, not all of the musicians were professional musicians. Similarly, athletes were categorized as recreational-athletes and no semi-professional, professional, or elite athletes were identified in the study. Distinguishing between multiple within-group levels may yield further findings. Furthermore, both expertise groups only participated in learning from object-label-ordering. Given the overall advantage of musicians in auditory processing (Chan et al., 1998; Cohen et al., 2011; George & Coch, 2011; Ho et al., 2003; Paraskevopoulos et al., 2017) and the increased salience and accuracy of short-term recognition memory of verbal over nonverbal stimuli (Calignano et al., 2021; Lukics & Lukács, 2022; Renvall et al., 2021; Talamini et al., 2022), musicians may have provided further evidence of general advantages of salient stimulus ordering in auditory-sequential associative word learning.

Finally, we will discuss the relative difficulty of associative word learning in the auditory domain as compared to the visual domain. In Cosper and colleagues (2022), visual behavioral performance accuracy was over 80%, while the auditory experiments yielded either no learning success (auditory sequential) or success rates below 70% (auditory simultaneous 66%). This seems to indicate that learning words for sounds in an associative manner is more difficult compared to learning words for visual objects. Beside the advantages of multimodal learning over unimodal learning and the integration of modalities in a multimodal language system (e.g., Eördegh et al., 2019; Matusz et al., 2015; Vigliocco et al., 2014), one potential further explanation for the visual advantage in relation to the specific type of stimuli used in our studies might be that the two stimuli clearly originate from different sound sources: environmental sounds vs. human speech. Auditory sequential processing does not only serve to discover links between all stimuli that naturally occur, but also to determine the sound sources of acoustic streams in so-called auditory scene analysis (Bregman, 1990; Winkler & Schröger, 2015). This might potentially be a more general problem in unimodal learning contexts, namely that the requirement to link two sounds competes with the necessity to segregate two sounds and assign them to different sources. Together with the general human preference for visual stimuli in adulthood, this might contribute to the relative difficulty of learning words for sounds, also reflected in the lower frequency of sound-related words compared to visual words (cf. Chedid et al., 2019; Chen et al., 2019; Lynott et al., 2020; Majid et al., 2018; Miklashevsky, 2018; Speed & Brybaert, 2022; Vergallito et al., 2020).

## 5 Conclusion

In the current study, we have shown that adult musicians do not have a behavioral advantage over non-musicians and athletes in auditory-sequential associative word learning. However, musicians do display evidence of implicit learning, albeit the learning may not be the result of lexico-semantic relationships. Furthermore, there is a behavioral advantage and the indication of a possible electrophysiological effect for label-object-ordering over object-label-ordering in auditory-sequential associative word learning. Our findings show how musical expertise and the salience of auditory-verbal stimuli influence association of sounds and labels. Future investigations into modality-specific associative learning should consider individual expertise as a factor in learning as well as saliency and memorizability of the stimuli in a given modality.

## Data Availability Statement

The data and scripts used for the analysis can be found here: https://osf.io/8wk9d/ and the experiments were not preregistered.

## Author contributions

**Samuel H. Cosper:** Conceptualization, Formal analysis, Investigation, Data Curation, Writing – Original draft, Visualization. **Claudia Männel:** Conceptualization, Writing – Review & editing, Supervision. **Jutta L. Mueller:** Conceptualization, Writing – Review & editing, Supervision.

## Acknowledgements

The authors would like to thank N. Gockel, L. Steinbach, C. Dammeyer, and all students and student research assistants of the Kindersprachlabor for their help in experiment coding and data collection. We are especially thankful to all of the participants.

